# Targeting glucose metabolism sensitizes pancreatic cancer to MEK inhibition

**DOI:** 10.1101/2021.01.09.425923

**Authors:** Liang Yan, Bo Tu, Jun Yao, Jing Gong, Alessandro Carugo, Christopher A. Bristow, Qiuyun Wang, Cihui Zhu, Bingbing Dai, Ya’an Kang, Leng Han, Ningping Feng, Yanqing Jin, Jason Fleming, Timothy P. Heffernan, Wantong Yao, Haoqiang Ying

## Abstract

Pancreatic ductal adenocarcinoma (PDAC) is almost universally lethal. A critical unmet need exists to explore essential susceptibilities in PDAC and identify druggable targets for tumor maintenance. This is especially challenging in the context of PDAC, in which activating mutations of KRAS oncogene (KRAS*) dominate the genetic landscape. By using an inducible *Kras^G12D^*-driven *p53* deficient PDAC mouse model (iKras model), we demonstrate that RAF-MEK-MAPK signaling is the major effector for oncogenic Kras-mediated tumor maintenance. However, MEK inhibition has minimal therapeutic effect as single agent for PDAC both in vitro and in vivo. Although MEK inhibition partially downregulates the transcription of glycolysis genes, it surprisingly fails to suppress the glycolysis flux in PDAC cell, which is a major metabolism effector of oncogenic KRAS. Accordingly, In vivo genetic screen identified multiple glycolysis genes as potential targets that may sensitize tumor cells to MAPK inhibition. Furthermore, inhibition of glucose metabolism with low dose 2-deoxyglucose (2DG) in combination with MEK inhibitor dramatically induces apoptosis in Kras^G12D^-driven PDAC cell in vitro, inhibits xenograft tumor growth and prolongs the overall survival of genetically engineered mouse PDAC model. Molecular and metabolism analyses indicate that co-targeting glycolysis and MAPK signaling results in apoptosis via induction of lethal ER stress. Together, our work suggests that combinatory inhibition of glycolysis and MAPK pathway may serve as an alternative approach to target KRAS-driven PDAC.

## Introduction

Pancreatic ductal adenocarcinoma (PDAC) is a highly aggressive malignancy with an overall 5-year survival of <10% and projected to be the second leading cause of cancer-related death by 2030 in the United States (1,2). PDAC is almost universally driven by mutationally activated KRAS, which represents the earliest and the most frequent (>90%) genetic alteration. However, no effective inhibitors have been developed for mutant KRAS due to the lack of suitable pockets except for small molecule inhibitors targeting KRAS^G12C^, which is present in just 1.5% human PDAC (3–5). To date, no targeted therapy has shown any major effect in PDAC patients yet.

KRAS mutation activates several downstream signaling pathways, including, but not limited to, the RAF/MEK/MAPK, phosphatidylinositol 3-kinase (PI3K)/AKT, and RalGDS pathways (6). Consequently, multiple cellular processes are activated such as proliferation, survival and KRAS-dependent metabolism pathways (7,8). However, not all of the pathways are activated by oncogenic KRAS simultaneously in any given tumor types. It is likely that a subset of KRAS surrogates play dominant roles during tumor maintenance. While several studies have established the roles of KRAS surrogates during PDAC initiation and development (5,9,10), the requirement of these pathways for tumor maintenance has not been thoroughly investigated. Recently, study using genetically engineered mouse (GEM) models indicates that C-RAF is required for the maintenance of Kras-driven lung adenocarcinomas (11).

Therapeutically, single agent targeting the MAPK or PI3K pathway has shown limited effects in PDAC (12,13). Although co-targeting MEK and PI3K pathways showed beneficial effect in preclinical models, the combination is too toxic to be used in clinic, indicating the need to identify alternative combination strategies. By using a PDAC GEM model driven by inducible *Kras^G12D^* (iKras model), we have established the critical roles of metabolism reprogramming in advanced tumors (14). Interestingly, recent studies have demonstrated the synergy between targeting metabolism pathways such as autophagy (15,16) or nucleoside metabolism (17) with MAPK inhibition in PDAC treatment, underscoring the therapeutic potential of co-targeting KRAS signaling and metabolism programs.

In this study, we conducted genetic studies to establish the dominant role of RAF-MAPK signaling, but not PI3K or RalGDS pathways, for KRAS-dependent PDAC maintenance. However, MAPK inhibition alone fails to suppress the glycolysis flux induced by oncogenic KRAS. Accordingly, genetic or pharmacological inhibition of glycolysis synergizes with MAPK inhibition to suppress PDAC growth both in vivo and in vitro. Moreover, molecular studies revealed that the synergistic effect is largely due to the induction of lethal ER stress and apoptosis. Our study identified the co-inhibition of MAPK signaling and glycolysis flux as an alternative approach to target KRAS-driven PDAC.

## Material and methods

### Animals

All animal manipulations were approved by the Animal Care and Use Committees at The University of Texas, MD Anderson Cancer Center under protocol number 00001549. No patient samples were directly used in this study. *TetO_Lox-Stop-Lox-Kras^G12D^ (tetO_LKras^G12D^), ROSA26-LSL-rtTA-IRES-GFP (ROSA_rtTA), p48-Cre* and *Trp53^L^* strains were described previously (18). Mice were fed with doxy water (Dox 2g/L in sucrose 20g/L) starting at the 3-week-old to induce PDAC development. For drug treatment, 10-week-old mice were treated by oral gavage delivery of Trametinib (1mg/kg/day), by intraperitoneal injection of 2DG (1000mg/kg/d) or both drugs together.

### Cell culture and reagents

IMR90 and human PDAC cell lines HPAC, 8988T, PaTu8902, Miapaca2, DanG, S2013, and PANC1 were purchased from American Type Culture Collection (ATCC). PDX148 was established from PDX tumors (19). IMR90 was grown in Eagle’s Minimum Essential Medium with 10% FBS. All human PDAC cell lines and PDX148 cells were cultured in RPMI1640 supplemented with 10% FBS. Mouse PDAC cultures derived from the iKras/p53 and LSL-Kras/p53 models were cultured in RPMI1640 supplemented with 10% Tetracycline Negative FBS (Gemini Bio Products). Trametinib, SCH772984, BKM120, GDC-0623, gemcitabine, and paclitaxel were purchased from Selleckchem. 4-Phenylbutyric Acid (4-PBA) and 2DG were obtained from Sigma Aldrich.

### Plasmids and reagents

The lentiviral shRNA clones targeting mouse aldolase A and nontargeting shRNA control were obtained from Sigma Aldrich in the pLKO vector. The clone IDs and sequences for shRNA are listed in the Supplementary Table S2. All ORFs were cloned into pHAGE-IRES-GFP using pENTR™/D-TOPO™ Cloning Kit (Thermofisher Scientific). The virus package plasmids psPAX2 (Addgene plasmid 12260) and pMD2.G (Addgene plasmid 12259) were purchased from Addgene.

### Glucose consumption and lactate production by YSI

Cells were seeded in 12-well plate at ~30% confluence (blank wells without cells were used for a baseline reading of Glucose/lactate). Forty-eight hours later, the culture medium was collected from each well and spinned at 1000g for 5min at 4°C. 250 μl supernatant was transferred into 96-well plate and read on YSI (Agilent). The number of cells in the 12-well plate was counted for normalization.

### Analysis of oxygen consumption rate (OCR) and glycolytic rate (ECAR) by Seahorse XF Analyzers

The determination of OCR or ECAR values was performed on XF96 analyzers. The iKras cells were seeded at a density of 1×10^4^ each well in Seahorse XF96 Cell Culture Microplate. The cells were washed twice with PBS on the second day and treated as Dox ON, Dox OFF or with Dox ON/TRA. After 24 h treatment, the cells were washed and cultured in Agilent Seahorse XF Base Medium, and the OCR or ECAR values were detected with Seahorse XF Cell Mito Stress Test following the manufacturer’s instruction. After seahorse analysis, the cells in 96-well plate were fixed with 4% paraformaldehyde for 15min, stained with DAPI and counted on Operetta High-Content Imaging System. The OCR and ECAR were normalized to the cell number in each well.

### Quantitative RT-PCR

Total RNA was extracted by the Qiagen RNeasy kit following the manufacturer’s instruction. cDNA was generated by using the SuperScript IV First-Strand Synthesis System (Invitrogen). The qRT-PCR was performed with Fast SYBR Green Master Mix (Applied Biosystems) in 96-well format in StepOnePlus (Applied Biosystems). Relative expression of genes was calculated by 2^-ΔΔCT^ method and normalized to beta-actin expression. All the used primers are listed in Supplementary Table S3.

### In vivo shRNA screen

The construction of customized shRNA library and method for shRNA screen in vivo were previously described (20,21). Briefly, targeting sequences of shRNA were designed using a proprietary algorithm (Cellecta) and 10 shRNA targeting each gene were included in the library. The polled shRNA was cloned into the pRSI16 lentiviral vector by using chip-based oligonucleotide synthesis. The oligonucleotide corresponding to each shRNA was synthesized with a unique molecular barcode (18 nucleotides) for measuring representation by next-generation sequencing. To perform the shRNA screen, the lentivirus package was prepared in 293FT cells using the second-generation packaging plasmids psPAX2 and pMD2.G. The lentivirus was concentrated using ultracentrifuge at 23,000 rpm for 3 h and the transducing unit was determined. The lentivirus infection was performed at ~0.3 transducing unit/cell with 10 μg/ml polybrene. After puromycin selection (2 μg/ml) for 48 h, cells were trypsinized, pooled together and 10^6^ cells were washed with PBS and stored in −80°C as reference. The remaining cells were mixed with matrigel (1:1) and orthotopically injected at 10^6^ cells/mouse pancreas. The screen was conducted in triplicates and an in vivo coverage of 1000 cells/barcode was guaranteed. The tumors were collected 10 days post injection and stored at −80°C.

### Extraction of tumor DNA and NGS library preparation

The frozen tumors were minced to small pieces and suspended in buffer P1 (QIAGEN, 1 mL Buffer/100 mg tumor) supplemented with 100 μg/mL RNase A (Promega). The dissociation of the tumor performed in disposable gentleMACS M tubes (Miltenyi Biotech) with the gentleMACS dissociator (Miltenyi Biotec). The cell pellet was suspended in buffer P1/RNAse A and lysed by adding 1/20 volume of 10% SDS (Promega). After incubating for 20 min at room temperature, the lysates were passed 10-15 times through a 22-gauge syringe needle to shear the genomic DNA. Then the genomic DNA were extracted using the Phenol-Chloroform solution. The DNA pellet was finally dissolved over-night in UltraPure distilled water (Invitrogen) and DNA concentration was assessed by NanoDrop 2000 (Thermo Scientific). To prepare the NGS libraries, the barcodes were amplified from the equal amount of genomic DNA by 2 rounds of nested PCR. The primers are list in Supplementary Table S4. The required adapters for NGS were introduced in the second PCR reactions. Amplified PCR products were purified using QIAquick gel extraction kit (Qiagen) and quantified using High Sensitivity DNA Assay (Agilent Technologies) for the Agilent 2100 Bioanalyzer.

### Screen data analysis

The shRNA screen data was analyzed as described previously (20,21). Illumina base calls were processed using CASAVA (v.1.8.2), and resulting reads were processed using our in-house pipeline. Following filtration and library-size normalization, reads counts in Vehicle or TRA samples were compared to the reference and a Log2 fold change was calculated.

### TEM

Cells were cultured in 12-well plate and washed with PBS twice before fixed with a solution containing 3% glutaraldehyde and 2% paraformaldehyde in 0.1 M cacodylate buffer (pH 7.3). Samples were then washed in 0.1 M sodium cacodylate buffer, treated with 0.1% Millipore-filtered cacodylate buffered tannic acid, post-fixed with 1% buffered osmium and stained with 0.1% Millipore-filtered uranyl acetate. The samples were dehydrated in increasing concentrations of ethanol and then infiltrated and embedded in LX-112 medium. The samples were then polymerized in a 60°C oven for three days. Ultrathin sections were cut using a Leica Ultracut microtome (Leica, Deerfield, IL) and then stained with uranyl acetate and lead citrate in a Leica EM Stainer. The stained samples were examined in a JEM 1010 transmission electron microscope (JEOL USA, Inc., Peabody, MA) using an accelerating voltage of 80 kV. Digital images were obtained using an AMT imaging system (Advanced Microscopy Techniques Corp., Danvers, MA).

### Cell Viability Assay

Cells were plated in equal number in 24 well plates and treated with 2DG, Trametinib or combination. After 4 days’ treatment, cells were rinsed twice with PBS to eliminate the floating cells and stained by Crystal Violet Staining Solution (0.25% Crystal violet in 20% methanol) for 20 min. The staining solution was removed and cells were washed with water. Stained cells were dried at room temperature and scanned. To quantify the relative cell numbers, cells were destained with 10% Acetic acid and absorbance was measured at 595 nm at appropriate dilutions. The Bliss Score for the combination was calculated by online tools (https://synergyfinder.fimm.fi/).

### Annexin V-PE and 7-AAD apoptosis assay

Induction of cell apoptosis was detected by PE Annexin V Apoptosis Detection Kit I (BD Pharmingen™) following the manufacturer’s instruction. Briefly, cells were seeded into 12-well plate at a density of 20000 cells/well and treated with the indicated concentration of 2DG, Trametinib or combination for 48 h. Then all the cells in each well were collected in 15ml tube and were pelleted by 200 g for 4 min. The supernatant was removed and cells were washed with PBS once. The cells were resuspended in the 100 μl 1×staining buffer in which 5 μl antibody and 5 μl 7-AAD were added. After staining at room temperature for 15 min, the samples were analyzed on Gallios Flow Cytometer (Beckman Counter). The FACS data were analyzed with FlowJo.

### Xenograft

For orthotopic xenografts, 5×10^5^ cells suspended in 10 μl 50% Corning Matrigel Matrix (Corning)/Opti-MEM media were injected into the pancreas of NCr nude mice.

For Sub-Q xenografts, 1×10^6^ cells suspended in 100 μl Opti-MEM media were injected subcutaneously into the lower flank of NCr nude mice. Animals were fed with doxy water and treated with Trametinib (1mg/kg/d), 2DG (1000mg/kg/d) or combination. Tumor volumes and body weight were measured every three days starting from Day 4 postinjection and calculated using the formula (Volume =0.5×length×width^2^).

### RNA sequencing

For the tumor samples, the mice were randomized into three groups (Kras-ON/Vehicle, Kras-OFF/Vehicle, Kras ON/TRA) on the 7^th^ day after orthotopic injection. After 1 or 3 days of treatment, the tumors were collected for RNA extraction. For the cell samples, 5×10^5^ iKras cell were seeded in the 10cm dish and started the treatment with vehicle, 1 mM 2DG, 25 nM TRA, or combination for 24 h. The RNA samples were collected in the TRIzol™ Reagent (1ml/10cm^2^).

Total RNA was extracted using the Qiagen RNeasy kit following the manufacturer’s instruction. The RNA samples with a RIN score >8 were used in further analysis. RNA library preparation, sequencing, raw data processing, and quality control were performed by Advanced Technology Genomics Core at MD Anderson Cancer Center. Reads were mapped using Tophat and FPKM values were generated with Cufflinks. The software package LIMMA (Linear Models for Microarray Data) was applied to detect significantly differentially expressed genes using Benjamini-Hochberg adjusted p-values.

### Immunohistochemistry and Western Blot Analysis

Tissues were fixed in 10% formalin overnight and embedded in paraffin. Immunohistochemical analysis was performed as described (22). Antibodies used for immunohistochemistry and WB were listed in Supplementary Table S5 and Table S6.

### Statistical analysis

To assess variance differences across various test groups, the data were analyzed using multiple t-tests in GraphPad Prism. Other comparisons were performed by using the unpaired 2-tailed t-test. For all figures with error bars, data are presented as mean ± SD unless otherwise stated. Tumor volume and tumor-free survival results were analyzed using GraphPad Prism. The level of significance was set at for p-value<0.01(**) or p-value <0.05 (*) in all figures.

## Results

### Active MAPK pathway is essential for PDAC maintenance

Our previous study has established the essential role of oncogenic KRAS for PDAC maintenance (14). To dissect the respective contributions of KRAS downstream pathways in tumor maintenance, three effector domain missense mutants of KRAS, which preferentially activate RAF/MEK/ERK pathway, PI3K pathway or RalGDS respectively (23,24), were ectopically expressed in the primary tumor cells derived from the iKras model *(p48Cre; tetO_Kras^G12D^; Rosa_rtTA^L/+^; p^L/+^)* (Fig. 1A). Using GFP as a negative control, complete tumor regression was observed upon Kras^G12D^ inactivation following doxycycline (Dox) withdrawal in orthotopic xenografts, indicating iKras line may serve as a powerful model to dissect the respective contributions of Kras downstream pathways for tumor maintenance (Fig. 1B). Interestingly, the Kras^G12V/T35S^, which selectively activates the RAF/MAPK pathway (Fig. S1A), functions as potent as the endogenous *Kras^G12D^,* to maintain xenograft tumor growth following doxycycline withdrawal (Fig. 1B). In contrast, Kras^G12V/Y40C^ (activates PI3K) or Kras^G12V/E37G^ (activates RalGDS) (Fig. S1A) was less efficient to maintain the tumor growth upon extinction of the endogenous *Kras^G12D^*. In consistent with the in vivo observation, the induction of sphere formation by Kras^G12V/T35S^ in the absence of endogenous Kras^G12D^ is comparable with Kras^G12V^. However, the number of spheres was reduced by about 90% if Kras^G12V/Y40C^ or Kras^G12V/E37G^ was expressed in iKras cells in the absence of doxycycline (Fig. 1C-D). Therefore, our data implicates the dominant role of RAF/MAPK signaling for KRAS-driven PDAC maintenance.

**Figure 1.**
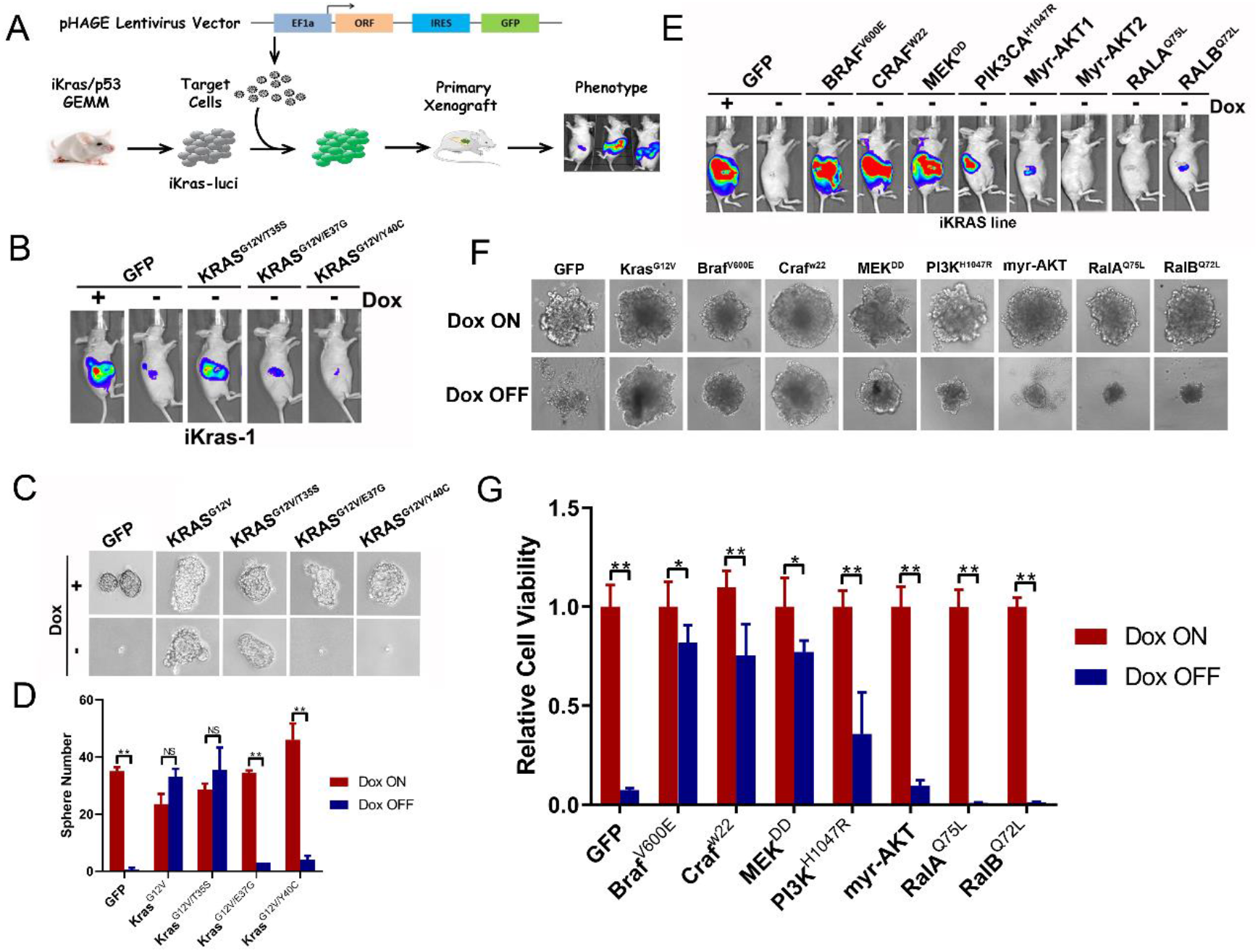
Active MAPK pathway is essential for PDAC maintenance. (A) Schematic diagram of investigating KRAS downstream surrogates in PDAC maintenance. The iKras cell was infected with lentivirus to overexpress the KRAS surrogates, sorted and orthotopically injected into nude mouse. Tumorigenesis was observed by bioluminescence imaging. (B) Tumorigenesis of iKras cells with Kras^G12V/T35S^, Kras^G12V/E37G^ or Kras^G12V/Y40C^ overexpression by bioluminescence imaging. (C) Sphere formation of iKras cell with Kras^G12V/T35S^, Kras^G12V/E37G^, Kras^G12V/Y40C^ overexpression in the low-attached plate. (D) Quantification of sphere number in C (n=3, Mean ± SD). (E) Tumorigenesis of mouse PDAC cells with constitutively active KRAS downstream surrogates by bioluminescence imaging. (F) Sphere formation of iKras PDAC cells with constitutively active KRAS downstream surrogates in low attached plate. (G) Quantification of spheres in F by cell viability assay (n=3, Mean ± SD).

To further dissect the KRAS effector pathways in advanced tumors, we conducted gain of function experiments using several constitutively activated mutants for the key effectors in MAPK, PI3K or RalGDS pathway (Fig. S1B-C). Consistent with our findings with the KRAS effector mutants, our data indicated that the constitutively activated C-Raf^W22^, BRAF^V600E^, and MEK^DD^ (25) were as efficient as mutant Kras to sustain tumor sphere formation in vitro and maintain tumor growth in vivo in the absence of doxycycline (Fig. 1E-G). In contrast, constitutive active PI3K^H1047R^ was less competent to support tumor sphere growth and induced much delayed tumor formation compared to the constitutive active mutants of the RAF/MAPK pathway components (Fig. 1F-G and S1D). Additionally, in the absence of doxycycline, iKras cells expressing Myr-AKT or constitutive active RalGDS (RalA^Q75L^ or RalB^Q72L^) failed to exhibit tumorigenic activity in vitro or in vivo (Fig. 1F-G and S1D). Together, our data indicates that RAF-MAPK signaling is the major pathway for KRAS-mediated PDAC maintenance.

### Inhibition of MAPK pathway fails to recapitulate glycolysis inhibition upon KRAS inactivation

Despite the essential role of MAPK pathway in PDAC maintenance, blocking MAPK signaling with MEK inhibitors has been shown to exert marginal impact on tumor growth or overall survival in both preclinical models and PDAC patients (12,13,26–28). Although the feedback activation of multiple RTKs and their downstream PI3K pathway has been shown to mediate the resistance to MPAK inhibition in KRAS-driven PDAC (12,13), co-targeting MAPK and PI3K is too toxic in PDAC patients despite the anti-tumor effect of this combination in preclinical model (12). In an effort to further explore potential MAPK co-inhibition targets, we leveraged the unique ability of the iKras model to genetically extinct oncogenic Kras in advanced tumors and conducted global transcriptomic analysis to identify KRAS downstream effector pathways/activities that are not affected by MAPK inhibition. Specifically, RNAseq analysis was performed in orthotopic xenograft tumors established with primary tumor lines from the iKras/p53 model to compare the transcriptomic profiles following KRAS extinction or MEK inhibitor (MEKi) treatment for 1 or 3 days (Fig. 2A-B).

**Figure 2.**
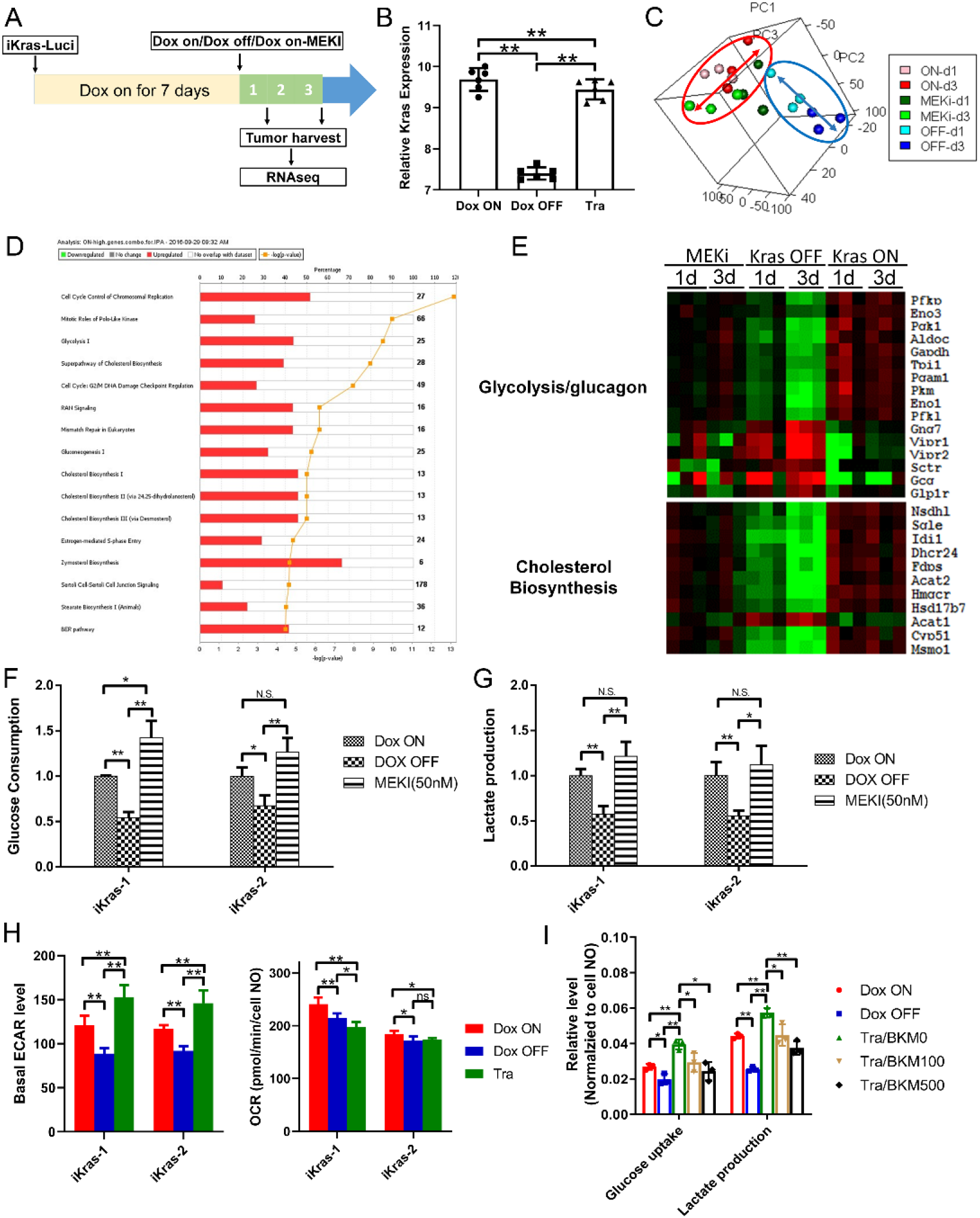
MEK inhibition fails to recapitulate glycolysis inhibition by Kras inactivation. (A) Schematic illustration of transcriptomic analysis between xenograft upon Dox ON, Dox OFF or TRA treatment in vivo. Seven days post orthotopic injection of iKras cells, the mice were randomized into three group (Dox ON, Dox OFF or treated with 1 mg/kg/day TRA) and xenograft tumors were collected after 1- or 3-days treatment. (B) Kras expression in the xenograft upon Dox ON, Dox OFF or TRA treatment by RNA-seq (n=5, Mean ± SD). (C) PCA analysis of transcriptome changes between Dox ON, Dox OFF or TRA treatment. (D) IPA pathway analysis of differentially expressed genes between Dox OFF and TRA treatment. (E) Heatmap of the representative differentially expressed genes in respective pathway. (F-G) Glucose consumption and lactate production of iKras cells with Dox ON, Dox OFF or TRA treatment by YSI (n=3, Mean ± SD). Expression validation of genes involved in glycolysis by qPCR (n=3, Mean ± SD). (H) Basal ECAR or OCR value of iKras cells with Dox ON, Dox OFF or TRA treatment by Seahorse (n=4, Mean ± SD). (I) Glucose and lactate concentration in the medium of iKras cell was measured upon treatment (TRA: 25nM; BKM120: 100 nM or 500 nM) for 48h. The glucose consumption and lactate production were normalized based on cell number (n=3, Mean ± SD).

The principal component analysis (PCA) revealed high concordance among the three biological replicates within each treatment group. Notably, a progressing shift was observed in Dox OFF 1-day, and 3-day tumors compared to ON Dox tumors, indicating the time-dependent transcriptomic change following KRAS extinction. However, such gradual expression changes were not observed in Trametinib-treated tumors (Fig. 2C), prompting the hypothesis that the pathways downstream of oncogenic KRAS that are not affected by treatment with MEKi could serve as coextinction targets that may cooperate with MEKi to recapitulate the impact of KRAS extinction. To this end, we conducted Ingenuity Pathway Analysis (IPA) on the differentially expressed genes in the Dox OFF group or the Trametinib treated group while compared with the Dox ON group. Half of the top-ten differentially expressed pathways preferentially enriched in OFF Dox tumors are associated with chromosol replication or cell cycle control (Fig. 2D and Fig. S2A), consistent with previous findings from an induced NRAS-driven melanoma mouse model (29). Interestingly, rest of the differentially enriched pathways are metabolism processes that we have previously shown to be driven by oncogenic KRAS in PDAC mouse models (14), including glucose metabolism and cholesterol biosynthesis (Fig. 2D). As shown in the clustered heatmap, compared to the dramatic downregulation of genes in glucose metabolism and cholesterol biosynthesis pathways, Trametinib treatment partially decreases the expression of these metabolism genes (Fig. 2E). In addition, Trametinib treatment also failed to recapitulate the induction of amino acid and fatty acid degradation pathways following KRAS inactivation (Fig. S2B).

Next, we employed different biochemical assays to determine the differential impact of KRAS extinction and MEK inhibition on glycolysis activity in mouse PDAC cells from the iKras/p53 model. Upon KRAS inactivation, glucose consumption or lactate production in the medium was dramatically downregulated, which is further supported by the decrease of extracellular acidification rate (ECAR) as measured with Seahorse. Surprisingly, no significant decrease in glucose consumption or lactate production was detected when the cells were treated with Trametinib for 2 days (Fig. 2F-G). Moreover, Seahorse analysis showed that Trametinib treatment lead to a mild induction of ECAR compared with the iKras/p53 tumor cells grown in the presence of doxycycline (Fig. 2H and Fig. S2C-D). Accordingly, MEK inhibition also failed to suppress the glucose uptake and lactate production in human PDAC cell lines (Fig. S2E-F). BKM120, a selective PI3K inhibitor, was able to reverse the AKT activation induced by Trametinib treatment and impaired the glucose consumption and lactate production (Fig. 2I and S2G), indicating that the sustaining of glycolysis activity upon MEK inhibition is PI3K-AKT-dependent in iKras cell. Together, our transcriptomic analysis and biochemical analysis indicate that MEK inhibition alone is not sufficient to suppress the KRAS-mediated glycolysis flux in PDAC cells, likely due to the feedback activation of PI3K signaling.

### Pooled shRNA library screening indicates glycolysis inhibition sensitizes iKras cells to MEK inhibition

To identify the metabolism genes that may sensitize Kras-driven PDAC cell to MEK inhibition upon depletion, we performed a pool-based in vivo loss-of-function screen in the orthotopic xenograft model as previously described (21). A customized bar-coded shRNA library comprising ~3,400 shRNAs targeting ~340 metabolism genes, including those KRAS-dependent metabolism genes (14) was packaged into lentivirus and infected iKras/p53 mouse PDAC cells. Orthotopic xenograft tumors were established in the nude mice and were treated with vehicle or Trametinib for 10 days before collection for next-generation sequencing. The log fold change (logFC) of bar-coded shRNA in the control or Trametinib-treated xenograft was calculated by comparison with reference group which was composed of library infected cells collected before orthotopic injection (Fig. 3A). In both control and Trametinib-treated xenograft, the positive control shRNA targeting PMSA1 and RPL30 are most depleted compared with the reference which indicates a high reliability of our screen system. Using logFC=-1 as a cut-off value, we obtained 36 candidate genes that were selectively depleted in MEKi-treated xenograft tumors compared to the control untreated ones (Supplementary Table S1). The depleted genes were most enriched in the glycolysis pathway (Fig. 3B), including Aldoa, Gapdhs, Hk2, Eno1 and PfkP (Fig. 3C-D).

**Figure 3.**
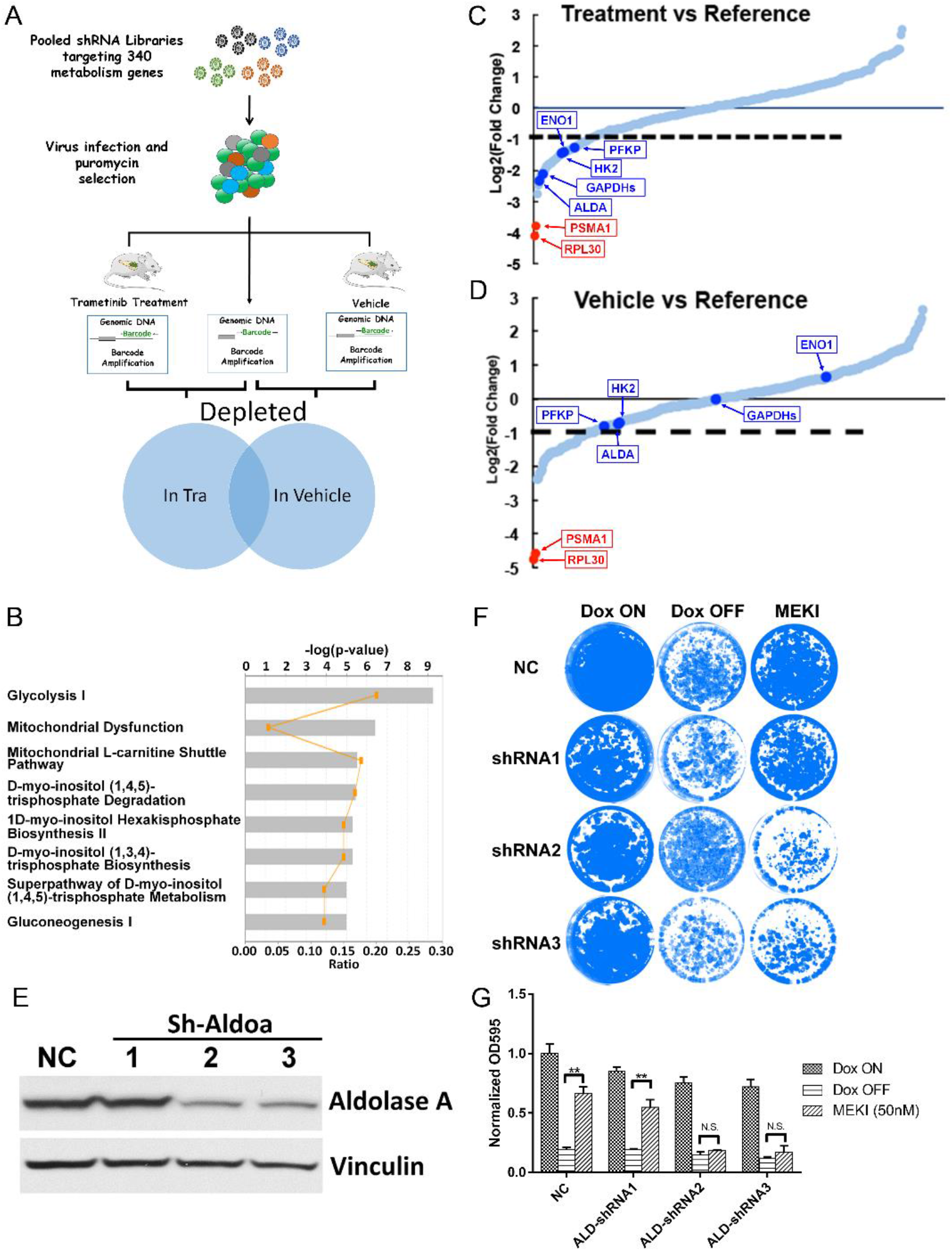
Pooled shRNA library screening indicates glycolysis inhibition sensitizes iKras cells to MEK inhibition. (A) Schematic illustration of pooled shRNA library screening in vivo. (B) IPA analysis of the depleted genes in shRNA library screen. (C-D) Relative abundance of each gene in the library in vehicle- or trametinib-treated tumors. Red dots: two positive control RPL30 and PSMA1; Blue Dots: five genes involved in glycolysis flux. (E) Knockdown of AldoA in iKras cell by shRNA. (F) Crystal violet staining of AldoA knockdown cell upon treatment of Dox ON, Dox OFF or 25 nM TRA for 4 days. (G) Quantification of crystal violet staining in F (n=3, Mean ± SD).

To validate the hits, we knocked down Aldoa in the iKras/p53 PDAC cells with shRNA (Fig. 3E). While knockdown of Aldoa itself has a limited effect on the cell growth, Aldoa-depleted cells are more sensitive to MEK inhibition compared with control cells, approximating the inhibitory effect on cell growth following KRAS inactivation (Fig. 3F-G). Together, our data suggests that glycolysis inhibition may sensitize the KRAS-driven PDAC cells to MEK inhibition.

### Glycolysis inhibition with 2DG synergizes with MEK inhibition to induce PDAC cell apoptosis

To test if pharmacological inhibition of glycolysis will synergize with MEK inhibition in KRAS-driven PDAC cells, we combined a well-known glycolysis inhibitor 2-deoxy-glucose (2DG) with Trametinib to treat iKras/p53 PDAC cells. While either 2DG or Trametinib treatment alone failed to significantly suppress cell growth, the combination dramatically decreased the proliferation of mouse PDAC cells from the iKras/p53 and LSL-Kras/p53 models, as well as human PDAC cell lines, including HPAC and Patu8902 cells (Fig. 4A-B). Interestingly, no synergistic effect was observed in normal lung fibroblast IMR90 (Fig. 4A-B), implicating a therapeutic window for such combination therapy. To evaluate the synergy over a broad range of 2DG and Trametinib concentration, we computed the Bliss independence score for Trametinib and 2DG combination in PDAC cell lines. The strong synergistic effect was indicated by a high positive score (=29.014) in the iKras/p53 PDAC cells but not in IMR90 cell (Bliss score=0.57) (Fig. 4C-D). The synergy between 2DG and Trametinib was also recapitulated in additional mouse LSL-Kras/p53 PDAC cells and human PDAC cells, including HPAC, and 8988T cells (Fig. 4E and Fig. S3A). Moreover, strong synergy was also observed between 2DG and additional MEK inhibitor GDC0623 or ERK inhibitor sch772984 (Fig. 4F and Fig. S3B). On the other hand, a low Bliss synergy score was observed when iKras cell line was treated with 2DG in combination with chemotherapy agents, such as gemcitabine or paclitaxel (Fig. 4G). Therefore, our data indicates that inhibition of glycolysis with 2DG specifically sensitizes PDAC cells to MAPK inhibition. Such synergy is due to the significant induction of apoptosis as shown by the increase in Annexin-V/7-AAD positive cells following the combination treatment, compared to single treatment alone groups. In contrast, the 2DG/Trametinib combination failed to induce apoptosis in IMR90 cells (Fig. 4H).

**Figure 4.**
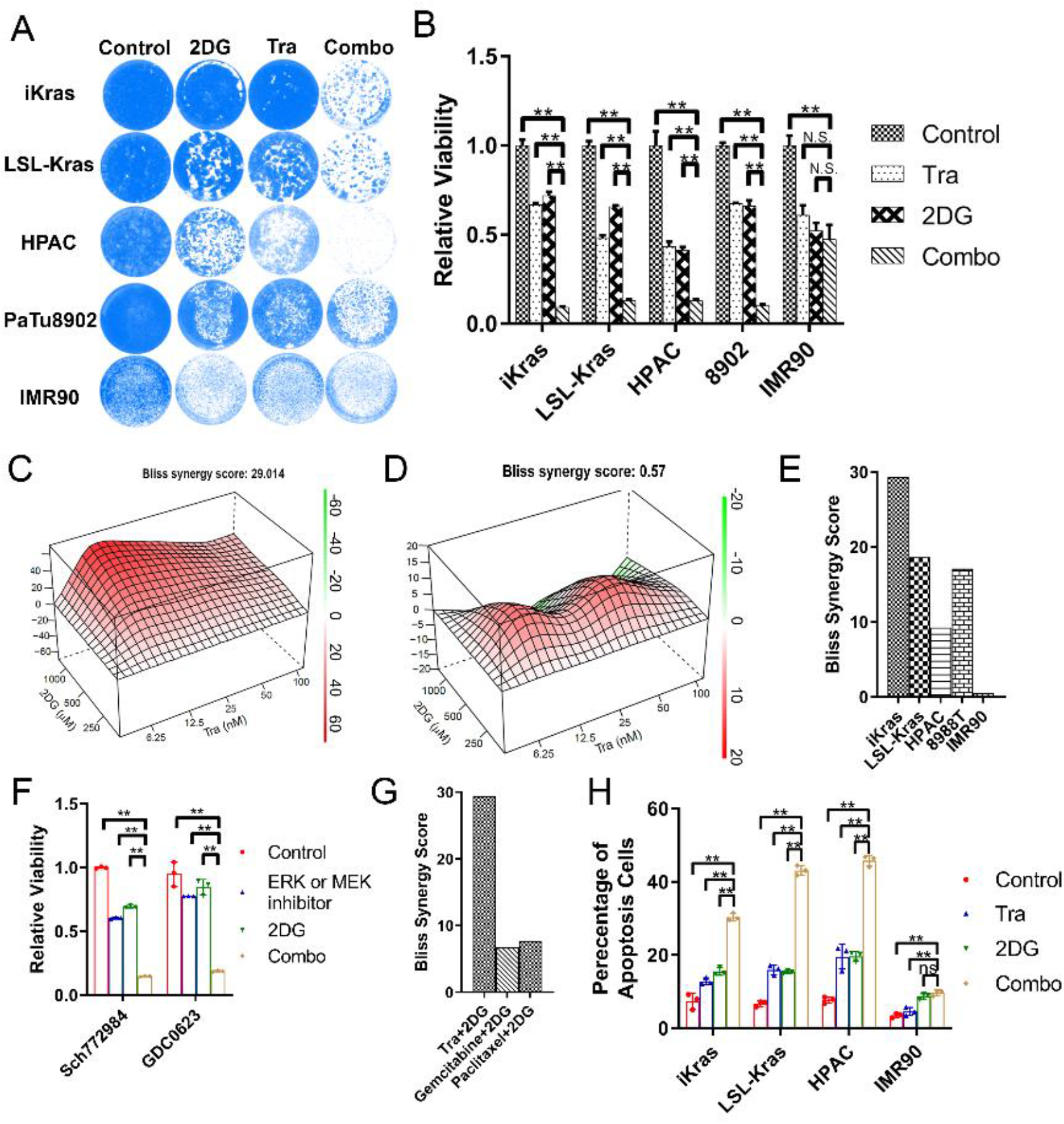
TRA and 2DG has a synergistic effect to induce apoptosis in mouse or human PDAC cell lines. (A) Crystal violet staining of the cells treated with 25nM TRA, 1mM 2DG or combination for 4 days. (B) Quantification of crystal violet staining in B (n=3, Mean ± SD). (C-D) Representative Bliss score for combination of TRA and 2DG in iKras or IMR90 cell. (E) The synergistic effect of TRA and 2DG combination was analyzed using Bliss score in mouse (iKras and LSL-Kras), human PDAC line (HPAC and 8988T) and IMR90 cell line. (F) Quantification of crystal violet staining for the cell treated with SCH772984 (200nM)/GDC063 (100nM), 2DG (1mM) or combination (n=3, Mean ± SD). (G) The Bliss synergy score was calculated for the combination of TRA/2DG, gemcitabine/2DG and paclitaxel/2DG in iKras cell. (H) Apoptosis of the cell treated with 25nM TRA, 1mM 2DG or combination by annexin V/7-AAD staining (n=3, Mean ± SD).

### 2DG and MEK inhibition synergistically induces apoptosis through ER stress

To gain molecular insight into the mechanisms underlying the synergy between 2DG and MAPK inhibition, transcriptomic analysis with RNAseq was conducted in the iKras/p53 tumor cells treated with Trametinib, 2DG or in combination. By using a cut-off of fold change>2 and p<0.01, a total of 850 up-regulated and 310 down-regulated genes were identified from three treatment groups compared to vehicle control group (Fig. S4A and S4B). IPA analysis of the 430 differentially expressed genes (308 up and 112 down) in the Trametinib group indicated the enrichment of signaling pathways such as the sirtuin signaling pathway (Fig. 5A). On the other hand, the 209 differentially expressed genes (177 up and 32 down) from the 2DG-treated group were enriched in metabolism pathways such as N-acetylglucosamine degradation (Fig. 5B). The largest number of differentially expressed genes were identified in the combination treatment group, including 526 up- and 210 down-regulated genes. The top two enriched pathways are autophagosome maturation and unfold protein response (UPR), which are not among the top enriched pathways in the single treatment groups, implicating the relevance to the apoptosis induced by the combination therapy (Fig. 5C).

**Figure 5.**
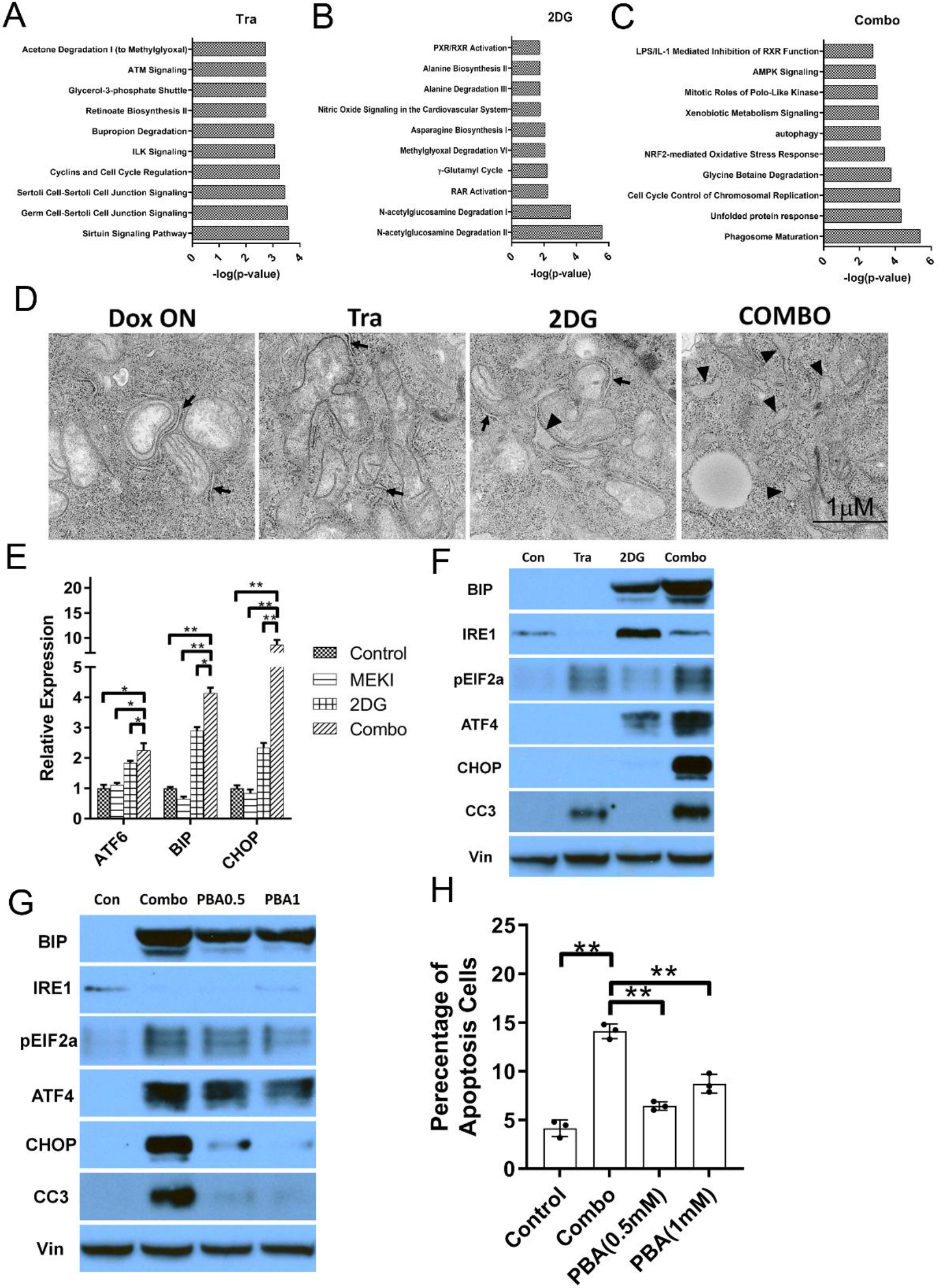
2DG and TRA has a synergy to induce ER stress in PDAC cell lines. (A-C) IPA analysis of differentially expressed genes in iKras lines treated with 2DG and TRA combination. (D) Representative TEM images of iKras cell treated with TRA, 2DG or Combination for 48h. The normal ER were indicated by arrow and swollen ER were indicates by arrowheads. (E-F) Expression of the ER stress markers in iKras treated with 25nM TRA, 1mM 2DG or combination for 48h detected by qPCR (E, n=3, Mean ± SD) or WB (F). (G) Expression of the ER stress markers in iKras treated with TRA/2DG or TRA/2DG/PBA by WB. (H) Apoptosis analysis of iKras cell treated with TRA/2DG or TRA/2DG/PBA by Annexin-V staining (n=3, Mean ± SD). TRA: 25nM; 2DG: 1mM and PBA: 0.5mM or 1mM.

Transmission Electron Microscope (TEM) revealed swollen ER in 2DG/Trametinib-treated iKras/p53 tumor cells (Fig. 5D), a morphology change indicating the induction of UPR. The activation of UPR was further supported by the drastic upregulation of ER stress markers, including ATF4, ATF6 and BIP, following 2DG/Trametinib treatment compared with vehicle control of single treatment groups (Fig. 5E-F). Moreover, CHOP, a well-known apoptosis activator downstream of UPR, along with cleaved caspase 3, was also specifically upregulated by the combination treatment in iKras/p53 tumor cells (Fig. 5E-F), indicating 2DG and MEK inhibition results in lethal ER Stress in PDAC cells. Interestingly, no induction of ER stress markers or cleaved caspase 3 was observed in 2DG/Trametinib-treated IMR90 cells (Fig. S4C). To further validate if unfold protein response is responsible for the induction of apoptosis by the combined 2DG and Trametinib treatment, cells were treated with the chemical chaperon PBA to decrease the UPR. As expected, PBA inhibited the induction of multiple ER-stress markers, such as BIP, ATF4, and phosphor-EIF2 α. Importantly, PBA treatment dramatically decreased the expression of CHOP, prevented caspase-3 cleavage and suppressed apoptosis induced by 2DG and Trametinib treatment (Fig. 5G-H), indicating the cell death induced by the combination treatment is mediated by hyper-activation of UPR.

### Trametinib in combination with 2DG exhibits antitumor activity in vivo

Next, we sought to evaluate the therapeutic potential of the combination in vivo. Briefly, the mouse iKras/p53 PDAC cells or PDX-derived human PDAC cells were injected subcutaneously into the immune-deficient mice. Tumor-bearing mice were treated with 2DG (1000mg/kg/d), Trametinib (1 mg/kg/d) or combination. Compared with single treatment group, combination group exhibited significant decrease in tumor size for both human and mouse PDAC (Fig. 6A-B). Moreover, we also evaluated the effect of the combination in the GEMM (iKras/p53^L/+^). After 7 weeks’ induction of KRAS^G12D^ expression with doxycycline, a time point previously showed to induce invasive carcinoma (14), the mice were randomized into four groups, including Vehicle control, Trametinib, 2DG, and Combo groups. While single treatment failed to elicit anti-tumor effect, combination treatment significantly prolonged overall survival (Fig. 6C). Immunohistochemistry revealed that the percentage of ki67 positive cells was significantly decreased in the combo-treated tumors compared to the single treatment groups, indicating inhibition of tumor cell proliferation (Fig. 6D and 6F). More importantly, BIP expression and percentage of cleaved Caspase-3 positive cells were significantly upregulated in the tumors treated with 2D/Trametinib combination (Fig. 6D and 6E), supporting the induction of UPR-related apoptosis.

**Figure 6.**
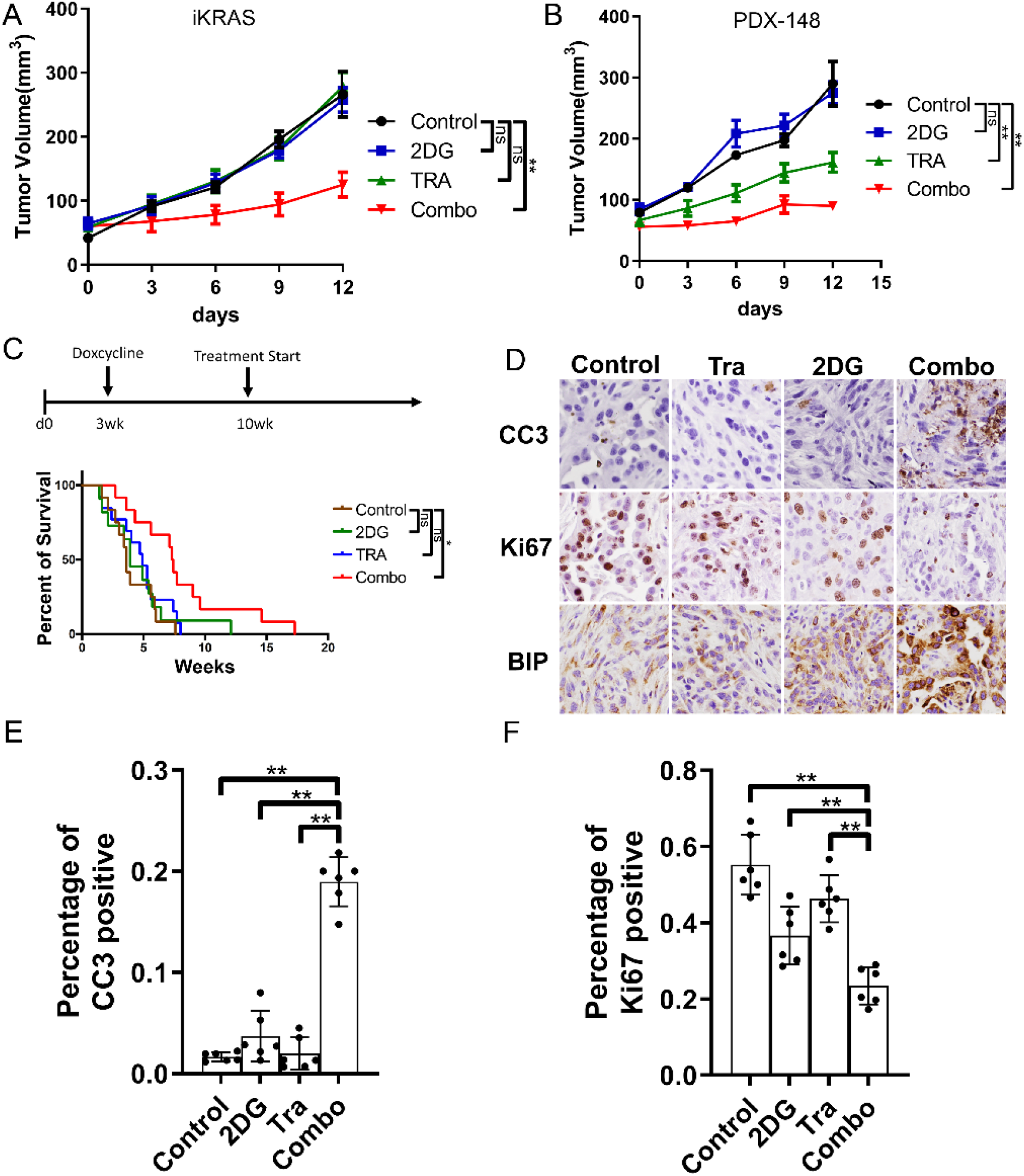
2DG and MEK inhibitor combination has a synergy to treat PDAC in vivo. (A) Xenograft tumor volume of iKras cell in nude mice treated with TRA, 2DG or Combo (n=5, Mean ± SE). TRA: 1mg/kg/day; 2DG: 1000mg/kg/d. (B) Xenograft tumor volume of PDX-148 cell in nude mice treated with TRA, 2DG or Combo (n=5, Mean ± SE). TRA: 1mg/kg/day; 2DG: 1000mg/kg/d. (C) Kaplan–Meier survival analysis of iKras GEMM treated with vehicle, TRA, 2DG or Combo (n=12). TRA: 1mg/kg/day; 2DG: 1000mg/kg/d. (D) Representative images of immunohistochemical staining of paraffin embedded xenograft tumors using antibodies for CC3, Ki67 or BIP. (E-F) Quantification of CC3 or ki67 positive cell in D (n=6, Mean ± SD).

## Discussion

As the major surrogates of KRAS signaling, the roles of MAPK and PI3K pathway in KRAS-driven tumors have been extensively studied. Mutations of BRAF, a dominant mediator for oncogenic KRAS signaling to activate MAPK signaling (30), were found to be mutually exclusive with the KRAS mutations in PDAC (5). Pancreatic-specific expression of oncogenic *Braf^V600E^* is sufficient to induce intraepithelial neoplasia (PanIN) lesions and invasive PDA in the autochthonous models (9). Moreover, CRAF has been shown to be essential for development of KRAS-driven non-small cell lung carcinoma (31,32), further supporting the central role of MAPK pathway in KRAS-driven tumorigenesis. On the other hand, PI3K and PDK1 have also been shown to be critical effectors downstream of oncogenic Kras in mediating cell plasticity and PDAC development (10). These data indicate that both MPAK and PI3K pathways are essential for KRAS-mediated tumor initiation. However, the requirement of these KRAS surrogates in advanced tumors has been less clear.

Recent study showed that ablation of CRAF expression leads to significant tumor regression in advanced tumors driven by KRAS^G12V^/Trp53 mutations (11), underscoring the requirement of MAPK pathway for tumor maintenance. Here our data provide additional evidences that MAPK pathway is necessary and sufficient for PDAC maintenance whereas PI3K activation is less competent to sustain tumor growth by itself, supporting the need to target MAPK pathway in KRAS-driven tumors. However, it has been well documented that targeting MAPK alone failed to elicit therapeutic benefit in KRAS-driven tumors, likely due to the feedback activation of PI3K pathway (12,33). Although co-targeting MAPK and PI3K was able to induce tumor regression and prolong survival in PDAC GEMM (12,13), the combination is too toxic to be tolerant in human patients (34). Previous studies have identified reactivation of multiple RTKs as a prominent mechanism of adaptive resistance to MEK inhibition in KRAS-driven tumors (13,33,35). However, co-targeting multiple RTKs is difficult to achieve therapeutically, pointing to the need to identify additional strategies to target PI3K or its downstream effector pathways.

In this study, we identified the potential role of PI3K-mediated glycolysis in the adaptive resistance to MEK inhibition in KRAS-driven PDAC. Co-targeting MAPK pathway and glycolysis with Trametinib and 2DG combination synergistically induces apoptosis in tumor cells both in vivo and in vitro. In line with our findings, recent study in BRAF-driven melanoma showed that glycolysis inhibitors were able to potentiate the effects of Braf inhibitor (36). Although MAPK signaling has been shown to mediate the transcription of multiple glycolysis genes downstream of oncogenic KRAS (14,17), here we showed that feedback activation of PI3K pathway is sufficient to maintain glycolysis flux in KRAS-driven tumors following MAPK inhibition. PI3K has been shown to be a master regulator for the transcription of glucose transporters (37). PI3K can also activate glycolysis at post-translational level by controlling the cytoskeleton remodeling and thus relieving the sequestration of glycolysis enzymes (38). Whether such mechanisms are also involved in the feedback activation of glycolysis upon MEK inhibition in KRAS-driven tumor cells remains to be elucidated.

Our data indicates that the maintenance of glycolysis activity is essential for the survival of PDAC cells following the inhibition of MAPK signaling. Blocking glycolysis with 2DG in combination with MAPK inhibition leads to induction of apoptosis. 2DG, a derivative of glucose, could be phosphorylated to 2DG-6-phosphate in cell. The accumulated 2DG-6-phosphate inhibits hexokinase in a noncompetitive manner (39) and can lead to the inhibition of multiple anabolic processes branched from glycolysis pathway. It’s possible that the synergy between 2DG and MAPK inhibition is due the blockade of multiple glucose-dependent metabolism pathways. Among them, 2DG has been shown to impair the pentose-phosphate pathway (PPP) and dramatically decreases R5P level (40). Interestingly, recent study has shown that the activation of PPP-mediated ribose metabolism is critical for the adaptation to the inhibition of KRAS signaling in PDAC cells (17). In addition, 2DG treatment has been shown to induce ER stress, likely due to its impact on HBP or mannose metabolism (41,42). While UPR is considered as a survival mechanism to maintain protein homeostasis, excessive ER stress will result in cell death (43,44). Here, we provide evidence that the combination of 2DG and MAPK inhibition dramatically amplified the ER stress in PDAC cells which results in apoptosis. This is supported by our finding that the cell death was partially rescued with chemical chaperon that mitigates the UPR. Therefore, our data indicates that the maintenance of protein homeostasis is critical for the survival of KRAS-driven PDAC cells upon the inhibition of MAPK signaling. It will be interesting to evaluate whether targeting additional regulators of protein homeostasis may also sensitize PDAC cells to MAPK inhibition.

Overall, our study provides evidence supporting the potential of co-targeting glycolysis and MAPK has an alternative approach to treat KRAS-driven PDAC.

## Acknowledgment

We thank the Institute for Applied Cancer Science for sharing reagents. We would like to thank Small Animal Imaging Facility, Histopathology Core, High Resolution Electron Microscopy Facility, the Flow Cytometry and Cellular Imaging Core at The University of Texas MD Anderson Cancer Center, and the Veterinary Medicine Department at MD Anderson (Cancer Center Support Grant CA016672). We thank Costas A. Lyssiotis for his suggestion and supports on the cell metabolism analysis. The research was supported by the National Cancer Institute (NCI) grant R01CA214793 and NCI P01 grant P01CA117969 to HY, and the Pancreatic Cancer Action Network-American Association for Cancer Research Pathway to Leadership Award to WY.

